# Post-interval EEG activity is related to task-goals in temporal discrimination

**DOI:** 10.1101/2020.08.18.256438

**Authors:** Fernanda D. Bueno, André M. Cravo

## Abstract

Studies investigating the neural mechanisms of time perception often measure brain activity while participants perform a temporal task. However, several of these studies are based exclusively on tasks in which time is relevant, making it hard to dissociate activity related to decisions about time from other task-related patterns. In the present study, human participants performed a temporal or color discrimination task of visual stimuli. Participants were informed which magnitude they would have to judge before or after presenting the two stimuli (S1 and S2) in different blocks. Our behavioral results showed, as expected, that performance was better when participants knew beforehand which magnitude they would judge. Electrophysiological data (EEG) was analyzed using Linear Discriminant Contrasts (LDC) and a Representational Similarity Analysis (RSA) approach to investigate whether and when information about time and color was encoded. During the presentation of S1, we did not find consistent differences in EEG activity as a function of the task. On the other hand, during S2, we found that temporal and color information was encoded in a task-relevant manner. Taken together, our results suggest that task goals strongly modulate decision-related information in EEG activity.

## Introduction

Perceptual timing is essential for humans and other animals to interact with their environments. A commonly used task to study this ability is temporal discrimination, in which participants have to judge whether a given duration is shorter or longer than a reference. Several studies have compared temporal discrimination tasks with discrimination of other attributes, such as color [1,2], size [3], space [4] and numerosity [5]. For example, Coull and colleagues [1], using functional Magnetic Resonance Imaging, found a higher activation of areas such as the pre-SMA and a network of other cortical and striatal areas when participants paid more attention to the duration than the color of a stimulus. In another study, Kulashekhar and colleagues [2] used a similar design combined with MEG to investigate possible neural correlates in temporal processing. Studies involving color discrimination as a contrast task change color dynamically to make the tasks more cognitively comparable. The rationale is that in both conditions, participants need to keep track of the visual stimulus presented. However, by changing color dynamically, time once again becomes relevant to the task: now, participants need to track for how long the target was presented with each hue. This limitation makes it hard to dissociate what aspects of neural activity are associated with temporal discrimination or with general task-related decisions in which time is relevant.

Here, we aimed to examine human electroencephalogram (EEG) using a duration and color discrimination task. In the color task, we used a static display so participants wouldn't use the temporal information to estimate the color. As in the study of Coull and colleagues [1], we controlled how much attention was allocated to different dimensions (time or color) by informing participants whether they would make a judgment about time or color before the stimuli (Pure Blocks) or only after the end of the trial (Mixed Blocks). Contrary to previous experiments, we parametrically varied the difference in time and color between stimuli, allowing an in-depth investigation of whether information about time or color in EEG activity was task-dependent. Thus, we aim to compare a simple interval discrimination task to a non-temporal task, in which time is not relevant to the decision.

Different time-resolved M/EEG markers have been proposed to be correlated with temporal judgments, such as the classical contingent negative variation (CNV) [6], the early post-interval N1P2 component [6], and the late positive component (LPC) [7–9]. This last marker has been studied in different explicit temporal paradigms, such as temporal bisection [7,8, 10], temporal generalization [7, 11], and temporal discrimination [3, 9, 12–14]. However, different studies have used the term LPC to refer to EEG activities diverse in time, topography, and task-related modulations. While some authors identified the LPC at prefrontal electrodes [3, 9, 11], others have placed it at centro-parietal electrodes [7, 10]. Here, we used a combination of multivariate methods of Linear Discriminant Contrast (LDC) [15] and Representational Similarity Analysis (RSA) [16] to evaluate dissimilarities of different activations of the time-resolved EEG signal in the different contexts of the experiment. With this method, it is not necessary to make a priori choices about electrodes and time points to analyze. Since the LDC is a dissimilarity measure across sensors, small changes in different electrodes will contribute to the final estimation, while choosing a set of electrodes for a simple event-related potential analysis might lose critical information.

In summary, we compared distances between patterns of EEG activity and investigated how these distances were modulated by task, by duration, or by color and whether possible modulations depended upon the task to be performed. We found weak differences between tasks when participants were exposed to the duration or color to be stored for further comparison. However, during the presentation of the comparison event, there were clear task-dependant differences in EEG activity. Decisions about durations and color evoked different patterns of EEG activity, at different moments and were modulated more strongly by task-relevant information.

## Materials and Methods

### Data availability

Task and analysis code and raw and pre-processed data are openly available (link to Open Science Framework project).

### Participants

Twenty-one human volunteers (age range, 21-32 years; 11 females) participated in the experiment. All of them had a normal or corrected-to-normal vision and did not report any psychological or neurological diagnoses. The Research Ethics Committee of the Federal University of ABC approved the experimental protocol (CAEE: 38370314.0.0000.5594), and the experiment was performed following the approved guidelines and regulations. Data from one volunteer (age 24, female, not included in the twenty-one participants above) were excluded from the analyses due to excessive noise and artifacts in the EEG signal (proportion of rejected trials above 20% in two segment windows of analyses as explained below).

### Stimuli and Procedures

The experiment consisted of a durationor color discrimination task (fig. 1). The stimuli were presented using Psychtoolbox [17] v.3.0 package for MATLAB on a 17-inch CRT monitor with a vertical refresh rate of 60 Hz, placed approximately in a viewing distance of 50 cm from the participant. Responses were collected via a response box of 9 buttons (DirectIN High-Speed Button; Empirisoft). We used the left and right buttons for responses in which participants should respond using both hands. We presented 720 trials consisting of two visual stimuli (filled circles) with different colors and durations. Participants were instructed to answer if the second stimulus was shorter/longer in duration or redder/bluer than the first one. The magnitude to judge was determined by the block condition: (1) In Time Pure and Color Pure blocks (2 blocks of each), participants were informed beforehand whether to judge differences in duration or color between the two visual stimuli; (2) In Mixed blocks (the remaining four blocks), participants would only know the magnitude (time or color) to judge during the response screen, 500ms after the offset of the second stimulus. Block order was randomized for all participants, and the background color was gray (RGB-color 100; 100; 100).

**Fig.1.**
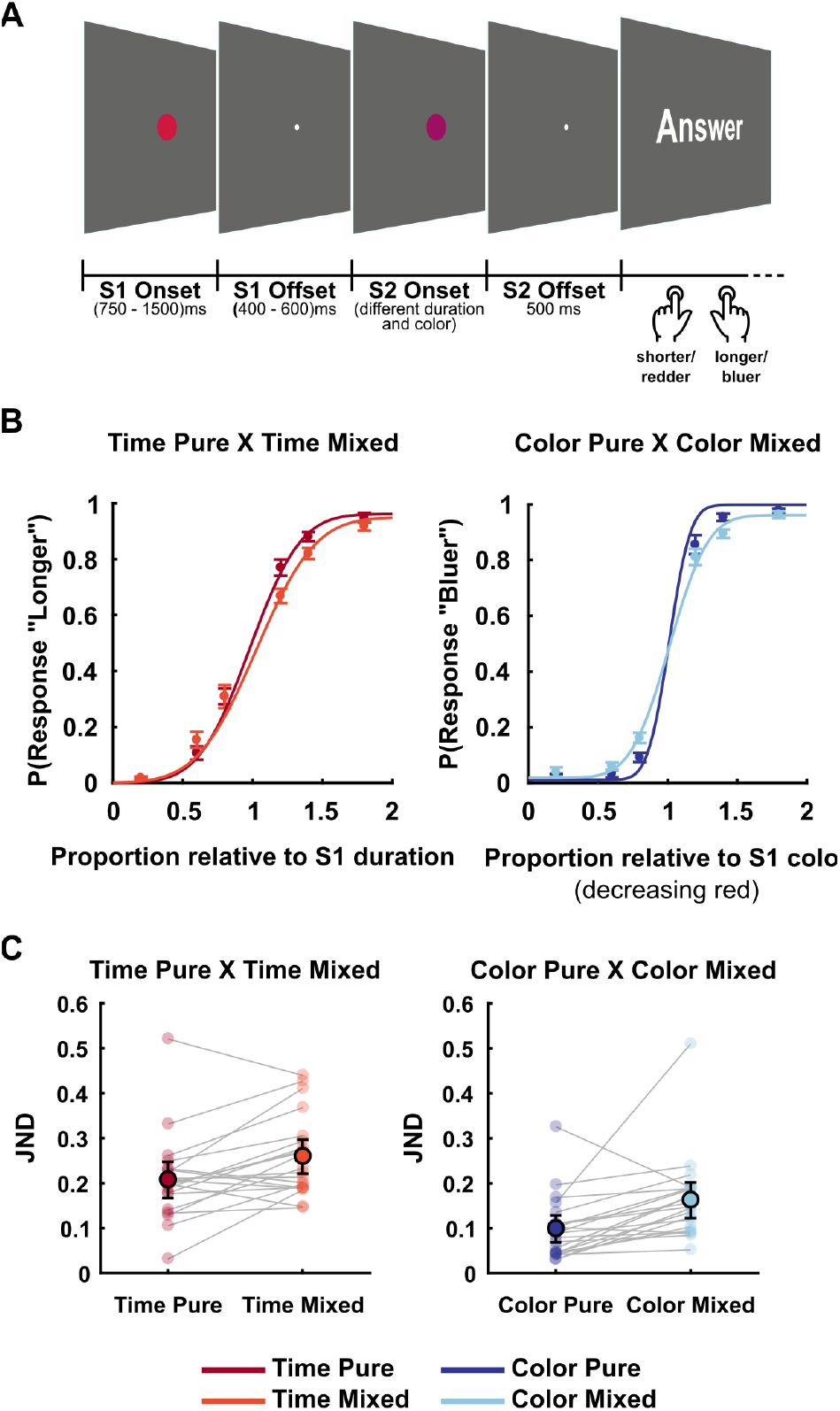
Experimental design and behavioral results. (A) Temporal/color discrimination task. The figure represents the time course of one trial. Before each block, a written cue would indicate if it was a ‘Time’ for Time Pure blocks, ‘Color’ for Color Pure blocks, or ‘Time/Color’ for Mixed blocks. (B) Curves show the psychometric functions for each condition, depicting the proportion of responding ‘longer’ for Time Pure and Time Mixed conditions (left) and responding ‘bluer’ for Color Pure and Color Mixed conditions (right). (C) Just Noticeable Difference (JND) by condition. Faint-colored filled circles represent individuals’ JND by condition. Lines connect JNDs for different conditions for each individual. Sharp-colored filled circles represent the mean JNDs, and bars represent the standard error of the mean.

Each trial started with the presentation of a circle (S1, one visual degree radius) at the center of the screen with a duration randomly chosen between 750 ms to 1500 ms, and colored in the RGB space [1-C, 0, C], in which C could range randomly from 0.2 to 0.5. The RGB space and parameter C for manipulating color was chosen based on [18]. After a random ISI of 400 ms to 600 ms, in which only a fixation point was present (0.25 visual degree radius), a second circle (S2, one visual degree radius) appeared with a different duration and color. Duration and color (controlled by parameter C) of S2 could range from 0.2 to 1.8 times the duration and color of S1 within6 possibilities in total: 0.2, 0.6, 0.8, 1.2, 1.4, 1.8. Durations and colors were independently randomized, and thus, orthogonal. After a delay of 500 ms, a response screen was presented in which participants were instructed to judge the duration or color of S2 relative to S1. In Pure blocks, the response screen reminded participants which dimension to be compared, while in Mixed blocks, the response screen informed which dimension should be compared.

### EEG recordings and pre-processing

EEG was recorded continuously from 64 ActiCap Electrodes (Brain Products) at 1000 Hz by a QuickAmp amplifier (Brain Products). All sites were referenced to FCz and grounded to AFz. The electrodes were positioned according to the International 10-10 system. Additional bipolar electrodes registered the electrooculogram (EOG). Data pre-processing was carried out using FieldTrip [19] toolbox for MATLAB. We segmented the data in four different epochs, for the period during the presentation of S1 and S2 (S1/S2 onset analysis) and just after the offset of each stimulus (S1/S2 offset analysis). Filters were applied to the continuous data with a bandpass of 0.1 Hz to 30 Hz (Butterworth filter, order 3). All data were re-referenced to the activity of electrodes TP9 and TP10, located in the earlobes and downsampled to 256Hz.

For the S1/S2 onset analysis, epochs were locked at the onset of S1/S2, and data were segmented from −150 ms to 750 ms. For the S1/S2 offset analysis, epochs were locked at the offset of S1/S2, and data were segmented from −150 ms to 400 ms for S1, and to 500 ms for S2. Channels with missing data due to problems in acquisition or channels with excessive noise were interpolated with neighbor channels using the FieldTrip channel repair function. Data from most participants had none or up to two channels interpolated. Only two participants had 3 and 4 channels interpolated.

For eye movement artifact rejection, an independent component analysis (ICA) was performed. Eye-related components were identified by the help of SASICA available for FieldTrip [20] and by visual inspection of topographies and time series from each component. Eye related components were then rejected for all segments. Baseline correction was performed using the periods from 150 ms before S1/S2 onset and 50 ms before and 50 ms after S1/S2 offset. Trials that exceed 200 μV for onset segments or 150 μV for offset segments were rejected. The percentage of rejected trials for S1 onset segment was 1.85% (range between 0% – 11.81%), for S2 onset was 1.18% (0% – 6.11%), for S1 offset was 0.75% (0%– 4.58%) and for S2 offset was 0.83% (0% – 4.58%).

### Behavioral Analysis

Behavioral analysis was based on the proportions of each type of response (longer/shorter or redder/bluer) as a function of the duration or proportional color of the second stimulus (S2) relative to the first stimulus (S1). We estimated psychometric functions for each participant in different conditions: Time Pure, Color Pure, and for the mixed blocks, we separated the data in trials in which participants were asked about duration (Time Mixed) and color (Color Mixed). Each of the four experimental conditions comprised 180 trials.

We fitted cumulative normal psychometric functions for each participant and condition, defined by four parameters: threshold, slope, lapse-rate, and guess-rate [21]. Guess rates and lapse rates were restricted to a maximum of 0.05. Each function's four parameters were fitted using maximum likelihood estimation as implemented in the Palamedes Toolbox [22]. To evaluate participants' performance, we estimated the Point of Subjective Equality (PSE) and the JND (Just Noticeable Difference). The JND is defined as the difference from 25% to 75% estimates of the psychometric curve, divided by two. This measurement represents how much different one stimulus has to be relative to another so that participants can notice. In contrast, the PSE represents the magnitude difference by which the second stimulus is equally likely to be judged as longer/shorter or redder/bluer than that of a first stimulus. We compared the JND and PSE from the Pure Blocks to their counterparts in the Mixed Blocks using a paired t-test. Effect sizes were estimated using Cohen's d as implemented in JASP [23].

### Multivariate Pattern Analysis

To compare the pattern of EEG activity across different conditions, we used Linear Discriminant Contrasts (LDC) [15]. We used this method to estimate distances in the time-resolved EEG signals from different tasks and conditions for each participant. All EEG electrodes were used in this analysis, except for the reference ones (TP9 and TP10). The LDC is a cross-validated Mahalanobis distance and allows the interpretation of ratios between distances, as its null distance is zero [15]. The LDC is calculated as:

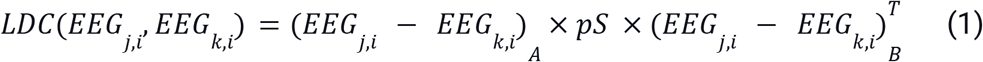

where *EEG_j,i_* and *EEG_k,i_* are row vectors of the means for each channel of the EEG activity in condition **k** and **j**, respectively, for each time point *ith*. *A* and *B* separate data in different subsets, representing different folds; *pS* is the pseudo inverse covariance matrix between *EEG_j,i_* residuals and *EEG_k,i_* residuals from subset *A*. We used a shrinkage estimator to calculate the pseudo-inverse covariance matrix *pS* [24–25]. Residuals were calculated by subtracting the activity of each trial, time point, and electrode from the mean activity for that electrode at that time point. The distance estimates are then averaged across all possible cross-validation folds.

Before estimating the LDC, data were smoothed within a 39 ms window. We used two-fold cross-validation to compare the electrophysiological activity during different experimental conditions. Folds were based using the blocked experimental design. For example, to compare EEG signals from pure conditions, we used each condition’s first block as one fold and the remaining block as the other fold. The analysis was conducted at each time point (3.9 ms apart after downsampling). To evaluate the estimated distances and correct for multiple comparisons across time, we used a mass-univariate approach. We used a permutation test over the *tmax* statistic, with strong control of the familywise error rate, as suggested by Groppe and colleagues [26]. All tests were one-sided t-tests compared with zero, and p-values were estimated using 10000 permutations. Significance values were based on an alpha level of 5% and we only considered significant windows ranging more than 20 ms.

To investigate how different aspects of time or color information influenced electrophysiological activity, we used a Representational Similarity Analysis approach. LDCs were calculated pairwise and used to create representational dissimilarity matrices (RDMs) for different intervals or color information by condition for each time point independently. We built theoretical matrices that represented the distances for time or color information for different comparisons separately. The resulting pairwise distances of the data and theoretical distances’ matrices were then entered into a simple linear regression analysis, separately by condition and segments of the experiment (S1 or S2, onset or offset) for each time point. The data-derived distances were dependent variables, and the theoretical distances matrices the independent variables. The estimated coefficients were compared to zero using a similar mass univariate approach as described above.

## Results

### Behavioral Results

The behavioral results (figure 1) showed that sensitivity, measured by the JND (Just Noticeable Difference), improved when participants knew beforehand which magnitude they would judge (Mean ± Standard error of the Mean, JND_TimePure_ = 0.207 ± 0.021, JND_TimeMixed_ = 0.259 ± 0.019, t(20) = −3.293, p = 0.004, d = 0.719; JND_ColorPure_ = 0.099 ± 0.015; JND_ColorMixed_ = 0.162 ± 0.020, t(20) = −3.309, p = 0.004, d = 0.722). There was no difference in bias, measured by the Point of Subjective Equality between mixed and pure blocks (PSE_TimePure_ = 0.961 ± 0.024, PSE_TimeMixed_ = 1.012 ± 0.030, t(20) = −2.070, p = 0.052, d = 0.452; PSE_ColorPure_ = 1.018 ± 0.022, PSE_ColorMixed_ = 1.016 ± 0.018, t(20) = 0.096, p = 0.924, d = 0.021). We assessed the goodness of fit using Tjur’s Coefficient of Determination [27] (mean D_TimePure_: 0.59, range 0.23 to 0.86; mean D_ColorPure_: 0.80, range 0.39 to 0.98; mean D_TimeMixed_: 0.50, range 0.29 to 0.72; mean D_ColorMixed_: 0.68, range 0.24 to 0.82). Our behavioral analyses showed that participants prioritized task-relevant information when they could anticipate the task to be performed.

### Electrophysiological Results

#### Exposure phase: No consistent differences in EEG activity by task-goals during S1

##### Task-related activity

In a first analysis, we focused on how activity evoked by S1 was modulated by the task to be executed. We aimed to investigate whether paying attention to the duration or the color of the stimuli leads to different stimulus encoding reflected in the EEG signal. Linear Discriminant Contrasts (LDC) were calculated comparing: i) pure conditions (Time Pure vs. Color Pure), ii) pure versus mixed conditions (Time Pure vs. Time Mixed and Color Pure vs. Color Mixed); and iii) between mixed conditions. The LDC was estimated from 150 ms before to 750ms after S1 onset (given that the shortest possible duration of S1 was 750 ms, this means that all not rejected trials were used). For S1 offset, we evaluated the EEG signal from 150 ms before S1 offset up to 400 ms (given that the shortest interval between S1 and S2 was 400 ms).

The mean distances between tasks during the exposure phase (S1) are shown in fig. 2A, and the spatial-temporal distribution of Event-Related Potentials (ERPs) for each task and period are shown in fig. 2B. Although there were short periods in which LDC exceeded the critical t-value (3.5) in Color Pure vs. Color Mixed conditions, distances between tasks were in general small and not significant. A similar pattern was observed for S1 offset, in which there were stronger distances between pure tasks, although not significant. Additional figures of topographies by tasks during S1 onset and at S1 offset can be found in the supplementary material at OSF (link to OSF).

**Fig. 2.**
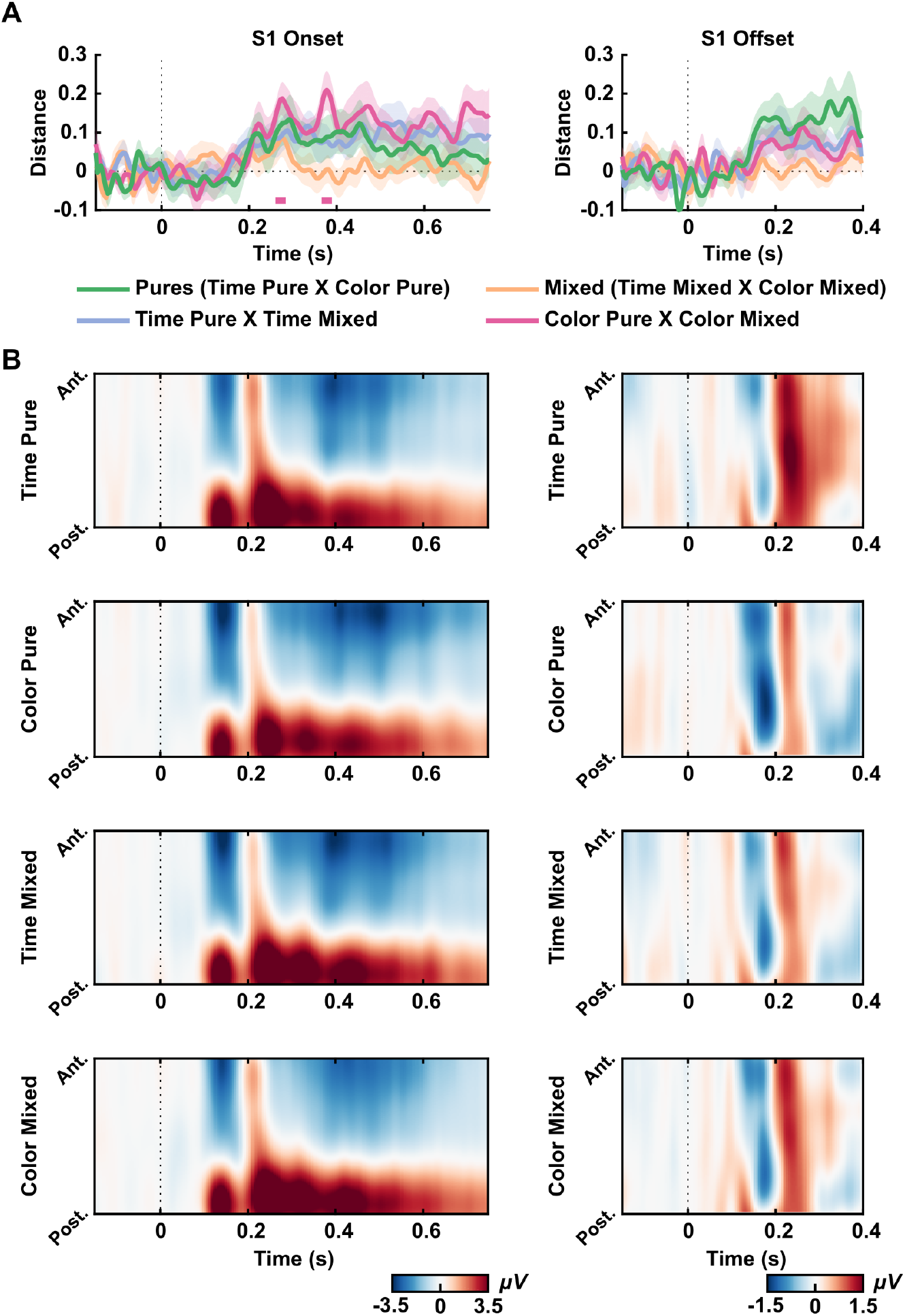
Exposure effects. (A) LDC distances between conditions for S1 onset (left) and S1 offset (right). The green curve indicates the mean distance between pure tasks (distance between Time Pure and Color Pure). The orange curve indicates the mean distance between mixed tasks (Time Mixed and Color Mixed). The purple curve indicates the mean distance between Time Pure condition and Time Mixed. The pink curve indicates the mean distance between Time Pure condition and Time Mixed. Shaded areas indicate Standard Error of the Mean. Straight lines indicate significant windows for distances from the permutation test. (B) Spatial-temporal ERPs for each condition from S1 onset (right) and S1 offset (left). Graphs show the mean electrical potential between participants from a topographical organization of electrodes (anterior to posterior) in time. All electrodes were used for plotting, except for the reference ones (TP9 and TP10).

##### Time and color-related activity

In a second analysis, we investigated if the duration or color of S1 modulated EEG activity. We aimed to test whether stimuli of different colors or durations evoked different patterns of EEG activity and whether this difference was more robust when that specific dimension was task-relevant. Color information was evaluated in the time-resolved S1 onset signal up to 750 ms. Given that information on how much time has passed since S1 onset was only available at S1 offset, we evaluated time information only at S1 offset. For each participant, S1 duration (from 750 ms to 1500 ms) or its color (indexed by the C parameter) were binned into six bins separately (around 30 trials for each time or color bins, for each experimental condition). The mean duration or mean C for each bin was used to calculate pairwise distances and build theoretical matrices. Pairwise LDC was calculated for each comparison. As explained in the methods section, these resulting pairwise LDC and the theoretical distances’ matrices were then entered into a linear regression analysis. In general, there were no consistent modulations of the EEG signal by color. For duration, we found one small period in which coefficients were larger than zero in Time Pure blocks (critical t = 3.5029; from 342.2 ms to 365.6 ms). The results can be seen in the Supplementary Figures at OSF (link to OSF).

#### Decisional phase: Consistent differences between tasks during S2

##### Task-related activity

In the next step, we focused on activity evoked by S2, the comparison stimulus. We performed the same LDC analysis to compare tasks during the second stimuli (S2 onset segments). To have a good number of trials of each condition and to have a considerable amount of time points during S2, this analysis was performed on data from trials in which the second stimulus lasted at least 750 ms.

For S2 onset, there was a consistent difference in the EEG signal between pure tasks (green line in fig. 3A left, window tested = −150 ms to 750 ms; critical t = 3.3922, significant distances from 185.9 ms to 748.4 ms). We also found differences for Time Pure and Time Mixed (purple line in fig. 3A left, critical t = 3.3722, from 455.5 ms to 486.7 ms, from 557.0 ms to 646.9 ms) and Color Pure to Color Mixed (pink line in fig. 3A left, critical t = 3.3942, from 232.8 ms to 271.9 ms, from 299.2 ms to 502.3 ms, from 510.2 ms to 557.0 ms, and from 572.7 ms to 748.4 ms). The spatial-temporal ERPs illustrate these differences measured by LDC from these conditions (fig. 3B, left). As expected, no significant distance was found between mixed tasks. In all significant comparisons, the difference was strongly driven by a centro-parietal p300 like response present in trials in which participants have to decide on the color of S2 (mixed blocks and pure color blocks). Additional figures of topographies by tasks during S2 onset and at S2 offset can be found in the supplementary material at OSF (link to OSF).

**Fig. 3.**
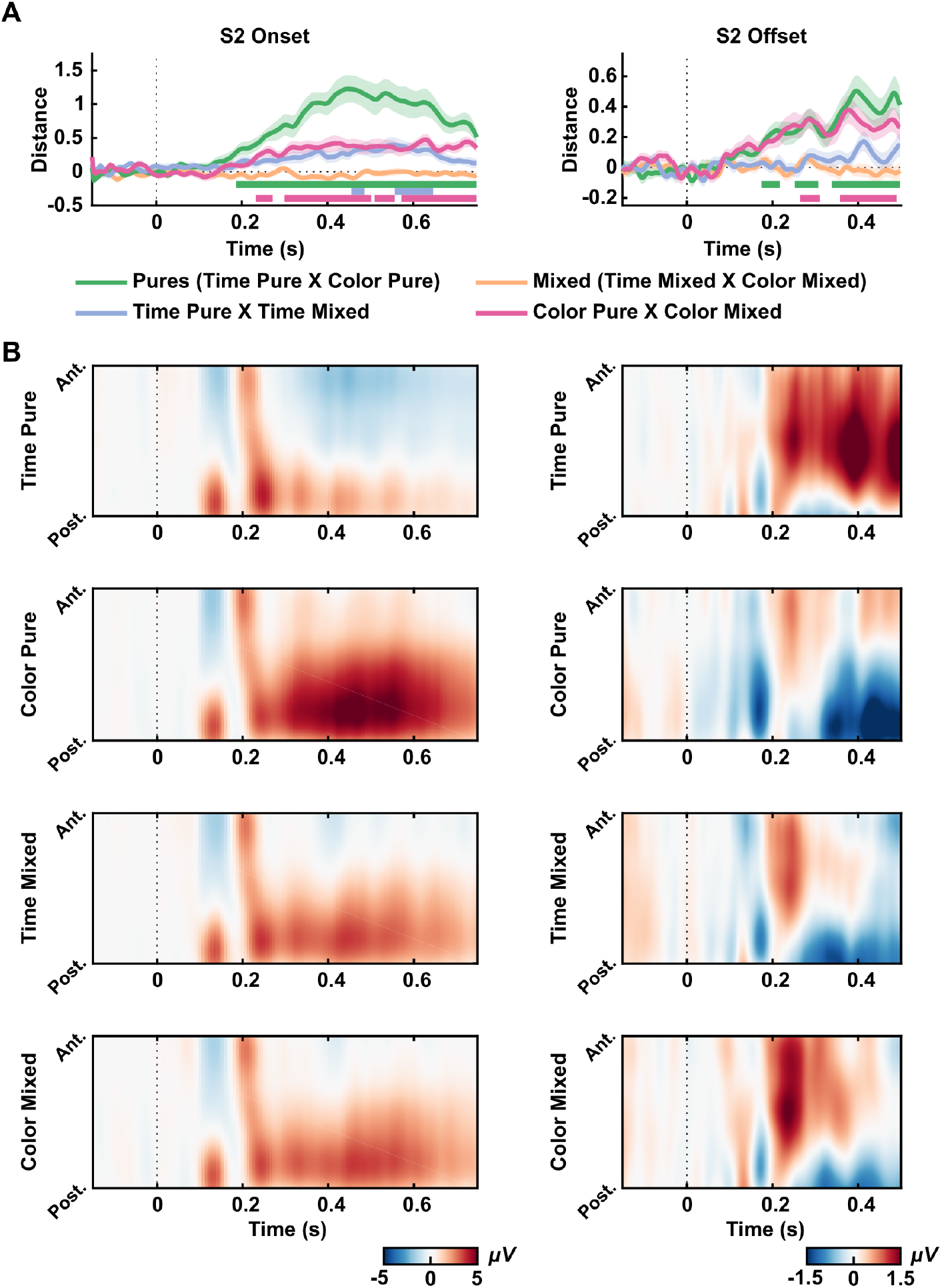
Decisional effects. (A) LDC distances between conditions for S2 onset (left) and S2 offset (right). Different colors indicate different comparisons. Shaded areas indicate Standard Error of the Mean. Straight lines indicate significant windows for distances from the permutation test. (B) Spatial-temporal ERPs for each condition from S2 onset (right) and S2 offset (left). Graphs show the mean electrical potential between participants from a topographical organization of electrodes (anterior to posterior) in time. All electrodes were used for plotting, except for the reference ones (TP9 and TP10).

The same analysis was conducted for S2 offset, from 150 ms before the offset of the stimulus to 500 ms after. Trials in which the second stimuli lasted less than 300 ms were excluded from this analysis to reduce sensory ERPs’ contamination. Again, there was a significant distance between pure tasks (green line in fig. 3A right, window tested = −150 ms to 500 ms; critical t = 3.3770, from 174.2 to 217.2 ms, from 252.3 to 307.0 ms and from 338.3 ms to 498.4 ms). There was also a significant distance between mixed and pure conditions (Color Mixed and Color Pure, pink line in figure 3A right, window tested = −150 ms to 500 ms; critical t = 3.3964, from 264.1 ms to 310.9 ms, and from 357.8 ms to 490.6 ms). Differences across conditions were strongly driven by EEG activity present in trials where participants have to decide on the duration of S2 (mixed blocks and pure time blocks). However, contrary to the p300 like activity, these differences seem to be more concentrated in frontal-central sensors.

#### Task-relevant features modulate post-interval EEG activity

##### Decision-related activity

We examined the modulation of EEG activity as a function of the stimulus magnitude of S2 relative to S1. An RSA approach was used to compare activity evoked by stimuli representing different proportions in time (proportional time) or color (proportional color) from S2 to S1, condition-wise. For time information, this analysis was performed on EEG activity of S2 offset (from 150 ms before the offset of the stimulus to 500 ms after) since full temporal information would be available only when the interval had elapsed. For color, this analysis was done for S2 onset up to 750 ms (for trials longer than this duration at S2), and we used the relative proportional color from S2 to S1. We calculated pairwise LDCs of the EEG signal for five possible time proportions of S2 relative to S1 (0.6, 0.8, 1.2, 1.4, 1.8), excluding the 0.2 proportion (see next) at S2 offset. Importantly, the 0.2 time proportion (in duration) was not included, given that this condition had only very short durations (maximum of 300 ms) to avoid false positives due to remaining evoked potentials from the onset and offset of the short duration visual stimulus. We also calculated pairwise LDCs of the EEG signal at S2 onset for all six possible color proportions (0.2, 0.6, 0.8, 1.2, 1.4, 1.8) for trials longer than 750 ms, thus we analysed the results up to this time point. These proportions were used to calculate pairwise distances and build two theoretical matrices, one relative to distances in time and one relative to distances in color (fig. 4A). As before, these matrices were entered into simple linear regression analyses, separately by condition, and coefficient estimates were evaluated.

**Fig. 4.**
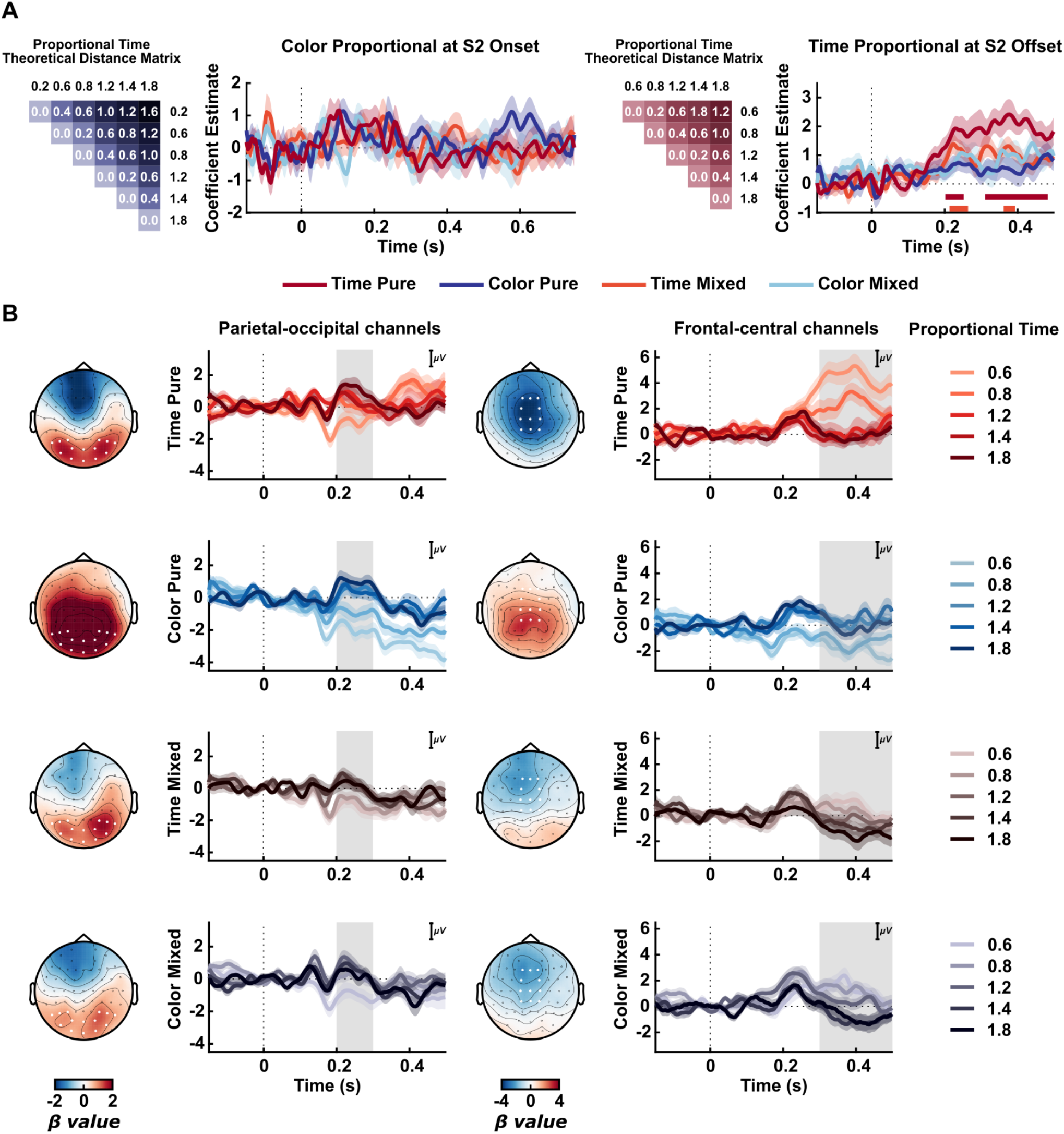
Proportional Time and Color Information at S2 offset. (A) Coeffcient estimates from RSA for relative proportional time information at S2 offset (left) and relative proportional color information at S2 onset (right). Matrices depict theoretical models used for RSA for time and color relative from S2 to S1. Curves show mean coeffcient values; shaded areas indicate Standard Error of the Mean. Straight Lines below indicate significant windows for RSA’s coeffcient estimates for each condition from the permutation test, represented by different colors. (B) Event-Related Potentials for different time proportions and conditions. Topographies of coeffcients (β) values from the mass univariate regression analysis are shown for different windows. The first column represents ERPs from the selected parietal-occipital electrodes (marked as white, P8, P6, P4, P3, P5, P7, PO8, PO4, POz, PO3, PO7, O1, Oz, O2) in the 200ms to 300ms window (gray area). The second column represents ERPs from the selected frontal-central electrodes (marked as white, F1, Fz, F2, FC1, FC2, C2, Cz, C1, CP1, CPz, CP2) in the 300ms to 500ms window (gray area).

As can be seen in fig. 4A (left column), we did not find a significant relation between color information and EEG activity at S2 onset (further exploratory ERPs figures are available on OSF).

However, for proportional time, we observed increasing coefficient estimates for time-relevant conditions (fig. 4A, right column). The RSA showed an increasing dissimilarity for proportional time information in the Time Pure condition (window tested = −150 ms to 500 ms; critical t = 3.3840, significant time windows from 201.6 to 252.3 ms, and from 310.9 ms to 482.8 ms), and in the Time Mixed (window tested = −150 ms to 500 ms; critical t = 3.4491, from 213.3 ms to 264.1 ms, and from 361.7 ms to 393.0 ms).

The modulation of EEG activity by proportional time information can be seen in fig. 4B. Based on the RSA analysis and the spatial-temporal evoked activity of S2 offset (fig. 3B, right column), we explored further ERPs at two different windows: from 200 ms to 300 ms and from 300 ms to 500 ms. For each of these periods, a linear regression between time proportions (0.6, 0.8, 1.2, 1.4, and 1.8) and EEG activity was performed for each participant and condition in each electrode and time point. As shown in these topographies (fig. 4B), these two intervals seem to illustrate two stages of the post-interval processing. A first parietal-occipital pattern (electrodes: P8, P6, P4, P3, P5, P7, PO8, PO4, POz, PO3, PO7, O1, Oz, O2) that has a positive correlation with time across all conditions. A second frontal-central pattern of activity (electrodes: F1, Fz, F2, FC1, FC2, C2, Cz, C1, CP1, CPz, CP2) shows a negative correlation between time and evoked activity: the shorter the second stimuli is from the comparison, the higher the amplitude of this late stage of this ERP.

To investigate further these two ERPs, we calculated the mean amplitude of the early activity at parietal-occipital electrodes and the late activity at frontal-central electrodes and estimated Spearman correlation coefficients separately per participant and task (fig. 5). At the group level, Spearman’s coefficients were Fisher transformed (replacing values of 1 and −1 to 0.95 and −0.95 before the transformation) and submitted to repeated-measures ANOVA. For the early parietal-occipital activity there were no significant differences of Spearman’s coefficients(ρ) across conditions (repeated measures ANOVA: F(3,20) = 1.011 p = 0.394, η² = 0.048).

**Fig. 5.**
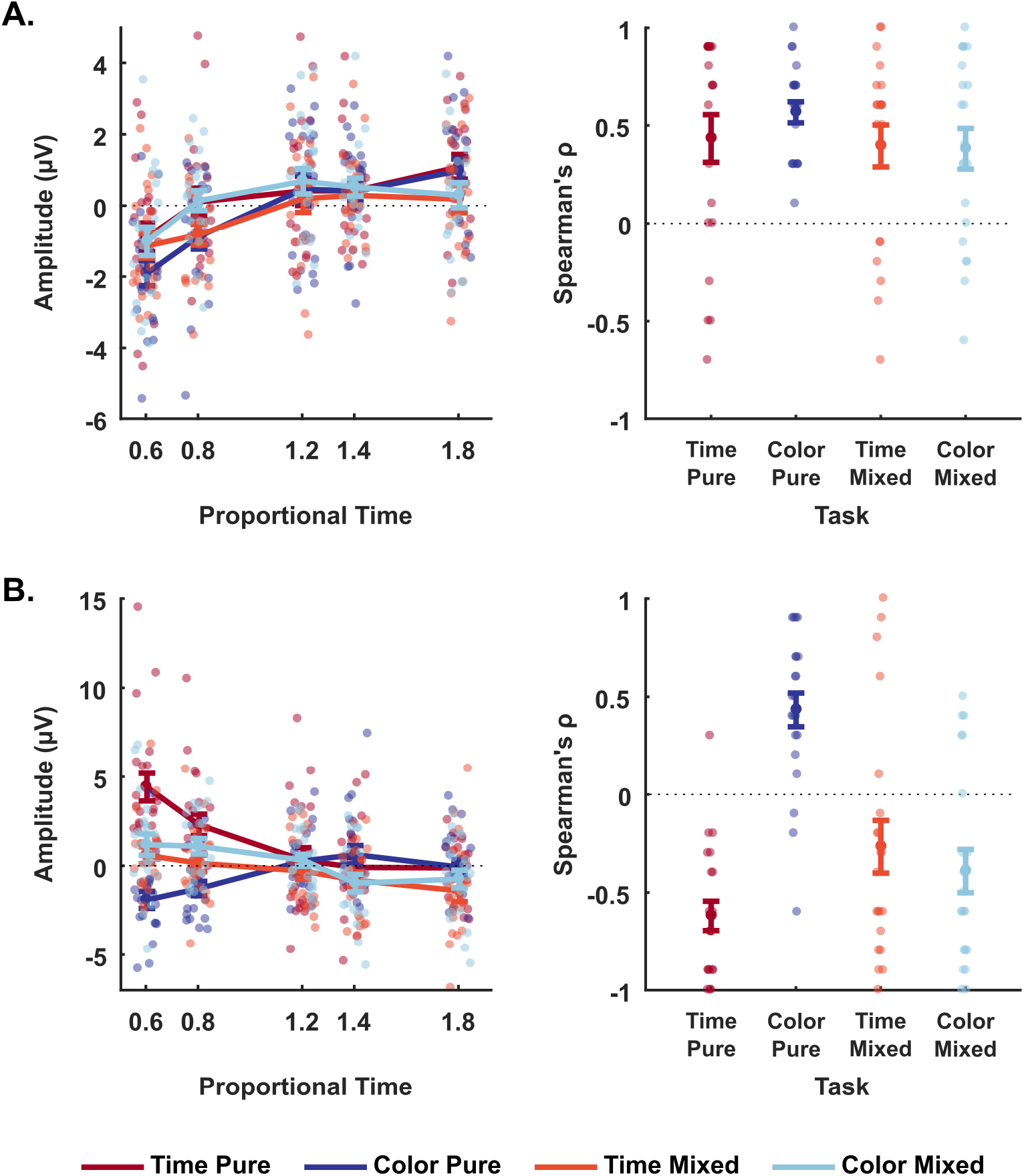
Correlation between proportional time and (A) early post-interval activity at the parietal-occipital electrodes, and (B) late post-interval activity at frontal-central electrodes. (A) Left: Mean amplitude at parietal-occipital electrodes (P8, P6, P4, P3, P5, P7, PO8, PO4, POz, PO3, PO7, O1, Oz, O2) in the 200ms to 300ms window, by proportional time and task. Right: Spearman’s coefficients (ρ). (B) Right: Mean amplitude at frontal-central electrodes (F1, Fz, F2, FC1, FC2, C2, Cz, C1, CP1, CPz, CP2) in the 300ms to 500ms window, by proportional time and task. Right: Spearman’s coefficients (ρ). Faint-colored circles depict individual values, while sharp-colored circles represent the mean between subjects. Bars represent the standard error of the mean.

For the late frontal-central activity, we found significant differences across conditions (F(3,20) = 30.116 p < 0.001, η² = = 0.601). Post-hoc comparisons (corrected using Holm–Bonferroni method) showed that the late frontal-central activity related to proportional time from the Time Pure condition is statistically different from the Color Pure (t(20) = −8.988, p < 0.001, d = −1.961) and the Time Mixed condition (t(20) = −2.818, p = 0.02, d = −0.615), but not from Color Mixed (t(20) = −1.964, p = 0.108, d = −0.429). Also, this late post interval activity related to time in the Color Pure condition is different from the two mixed conditions (Time Mixed: t(20) = 6.170, p < 0.001, d = 1.346; Color Mixed: t(20) = 7.023, p < 0.001, d = 1.533), but is not different across mixed conditions (t(20) = 0.854, pholm = 0.397, d= 0.186).

In general, these results suggest that the modulation of this frontal-central activity was strongly modulated by whether time was relevant or not to the current task.

#### The relation between post-interval activity, time and behavior is task-dependent

To evaluate whether there is a relation between the post-interval activity and the response given by participants, we performed a binomial regression in which the binary answer (shorter or longer) was used as the response variable and the proportional time (5 values: 0.6, 0.8, 1.2, 1.4 and 1.8) and EEG activity residuals were explanatory variables. We performed this regression for all conditions for the two different sets of electrodes and the two different windows after S2 offset (the same windows and electrodes explored at EEG activity topographies in figure 4B). We calculated EEG activity residuals by subtracting the mean EEG amplitude for that specific proportion of S2 and condition in each electrode and time point. The residuals of EEG activity were used to minimize the correlation between proportional time and EEG activity. This allowed us to investigate whether trial-by-trial EEG fluctuations covaried with behavior.

As expected, proportional time was a significant predictor of behavior for all time-relevant conditions (Time Pure condition: first windows: t(20) = 11.63, p < .0001, d = 2.54 and second windows: t(20) = 11.49, p < .0001, d = 2.51; Time Mixed condition: first windows: t(20) = 12.64, p < .0001, d = 2.76 and second windows: t(20) = 12.56, p < .0001, d = 2.74), but not for color relevant conditions (Color Pure condition: first windows: t(20) = −0.46, p = 0.65, d = −0.10; and second windows: t(20) = −0.48, p = 0.63, d = −0.11; Color Mixed condition: first windows: t(20) = 0.88, p = 0.39, d = 0.19 and second windows: t(20) = 0.86, p = 0.40, d = 0.19).

Critically, the early EEG post-interval activity modulated behavior only in the Time Mixed Condition (t(20) = 2.53, p = 0.02, d = 0.55), but not on other conditions (Color Mixed (t(20) = 1.55, p = 0.14, d = 0.34, Time Pure (t(20) = 1.18, p = 0.25, d = 0.26, Color Pure (t(20) = 0.41, p = 0.69, d = 0.09). On the other hand, late EEG post-interval activity was associated with temporal judgements for both Time Pure (t(20) = −2.97, p = 0.01, d = −0.65) and Time Mixed (t(20) = −2.34, p = 0.03, d = −0.51) but not for Color Mixed (t(20) = −1.67, p = 0.11, d = −0.36) nor Color Pure (t(20) = 0.46, p = 0.65, d = 0.10).

## Discussion

In the present study, we investigated the neural correlates of temporal discrimination. To dissociate duration-related EEG from other task-related decisions, we used multivariate analyses to compare activity across two tasks: one in which participants had to compare the color of two stimuli and one they had to compare their durations. Across different blocks, participants did or did not have prior knowledge about what feature would have to be compared. As expected, our behavioral results showed a better performance in conditions that participants knew the feature to be compared.

Our task consisted of the encoding of duration/color information of a first stimulus and a decision about these features in a second stimulus. Using an MVPA approach, we were able to investigate patterns of EEG activity, without needing to select moments and groups of sensors *a priori*, and compared whether and how EEG activity differed due to task-goals in these two phases. In general, we observed that: (1) During the encoding phase, EEG activity did not differ strongly across tasks; (2) During the decision phase, there were substantial differences in EEG activity across tasks and how task-relevant features modulated this activity.

Our task design allowed a temporal separation of when information about each feature was accessible to participants during the decision phase. While color information was available at the onset of S2, information about time was fully present only at its offset. This separation was reflected in the EEG signal, in which making decisions about color or time evoked activity at the onset and offset of S2, respectively. When making color decisions, there was a clear p300 at S2 onset, in agreement with proposals of this potential reflecting decision-making processes [28–29]. Although the p300 was not strongly modulated by how different S2 was relative to S1, this could be due to the color task being slightly easier for participants. On the other hand, decisions about durations evoked a more robust pattern of activity in fronto-central sensors, only at the offset of S2. Critically, this offset activity modulation by duration was most reliable when the temporal information was task-relevant, and weaker when it was irrelevant or only possibly relevant.

Our findings are in partial agreement with a by Kulashekhar et al. study [2] that did not find differences in the time-locked signal during the encoding period in MEG recordings. However, contrary to our findings, the authors did not find differences in the time-locked signal during the decision period. There were substantial differences between our tasks that might explain the contrast between these findings. In their work, the color task consisted of a varying hue that changed during the whole trial from a bluish to a reddish-purple. This was done as an attempt to make the color and the temporal task more comparable, given that participants would have to integrate information across the whole trial to make their color decision in a way similar to the temporal task [1–2]. However, even with this control, it was possible that participants could still accumulate enough color information during S2, although in different moments across different trials [2], as suggested by the difference in reaction times between tasks. Another crucial difference is that temporal information could be used in the color discrimination task for participants to estimate the mean color. When evaluating the predominant color, participants can calculate for how long each hue was presented. On our task, color discrimination task is not time-dependent, and thus the moment in which task-related information was made complete was clearer: at S2 onset for color and at S2 offset for time. This allowed a more direct comparison and showed that decisions about time and color evoked different activity patterns.

An increasing number of studies have suggested that EEG markers at the end of the interval are correlated with temporal processing, such as the early post-interval N1P2 component [6] and the late positive component of timing [7–9]. In agreement with these findings, we found a modulation of post intervals signals by time in EEG activity that resembled a parieto-occipital p200 and a later fronto-central similar to the LPC. However, only the LPC seemed to be more strongly correlated with time and behavior.

Our results corroborate with previous findings that the LPC might be related to decisional stages on other temporal tasks, such as temporal bisection [7,8, 10], temporal generalization [7, 11] and temporal discrimination [9,3, 12–14]. However, it is important to stress that different studies have used the term LPC to refer to EEG activities diverse in time, topography, and task-related modulations. While some authors have measured the LPC to the response [7, 8, 10], others have measured it relative to the offset of the interval [3, 9, 12–14]. Additionally, different authors have identified the LPC at prefrontal electrodes [3, 9, 11], and centro-parietal electrodes [7, 10].

In our results, the LPC had a fronto-central distribution and was inversely correlated with how much shorter the comparison interval was relative to the reference, in agreement with previous studies [7, 10]. This inverse relationship between time and LPC amplitude seemed to hold only for intervals shorter than the reference, while intervals longer than the reference had a similar amplitude. This pattern is consistent with the proposal that a decision only needs to be made at the offset of the interval when the duration is shorter than the reference [30].

Recent proposals have approximated temporal processing with drift-diffusion models of decision-making [31–33]. When adapted to a temporal discrimination task like ours, these models posit that at the offset of the interval to be judged, evidence accumulates towards one of two thresholds. Our findings are consistent with this proposal, with the pattern of accumulation reflected mainly on the LPC. Importantly, although the LPC had a temporal distribution similar to other activities commonly associated with a decision, such as the p300 and the centro-parietal positivity (CPP) [28, 29], its scalp topography was more fronto-central than the p300/CPP. Critically, as mentioned above, the LPC in our findings had a higher amplitude for intervals that are shorter than the reference, consistent with the proposal that this decision process should take place only for shorter than reference intervals [31–33].

In conclusion, our results suggest that task goals strongly modulate temporal information encoding in EEG activity. Future studies, using similar approaches, should investigate whether and how different temporal tasks modulate this activity pattern and whether it is present only in decisions about time, or in other forms of decisions that evolve monotonically in time.

## Acknowledgements

The authors would like Gustavo Rohenkohl and the Timing and Cognition lab for comments on an earlier version of the manuscript.

## Author contributions

Conceptualization: Fernanda Dantas Bueno, Andre Mascioli Cravo

Data curation: Fernanda Dantas Bueno

Formal analysis: Fernanda Dantas Bueno, Andre Mascioli Cravo

Funding Acquisition: Andre Mascioli Cravo

Investigation: Fernanda Dantas Bueno, Andre Mascioli Cravo

Methodology: Fernanda Dantas Bueno, Andre Mascioli Cravo

Project administration: Andre Mascioli Cravo

Resources: Andre Mascioli Cravo

Supervision: Andre Mascioli Cravo

Visualization: Fernanda Dantas Bueno

Writing – original draft: Fernanda Dantas Bueno, Andre Mascioli Cravo

Writing – review & editing: Fernanda Dantas Bueno, Andre Mascioli Cravo

## Conflict of interest statement

The authors declare no conflict of interest

## Data Availability Statement

All the files are available in https://osf.io/632e7/?view_only=a3a64f082f6f4f2f8ed0d4a6c60d2010

## Funding

F.D.B. was supported by grant 2017/24575-3, São Paulo Research Foundation (FAPESP). A.M.C. was supported by grant 2017/25161-8, São Paulo Research Foundation (FAPESP).

## Notes

### Competing Interest Statement

The authors have declared no competing interest.

## References

1. Coull, J. T., Vidal, F., Nazarian, B., & Macar, F. (2004). Functional anatomy of the attentional modulation of time estimation. Science, 303(5663), 1506–1508.

2. Kulashekhar, S., Pekkola, J., Palva, J. M., & Palva, S. (2016). The role of cortical beta oscillations in time estimation. Human brain mapping, 37(9), 3262–3281.

3. Gontier, E., Le Dantec, C., Paul, I., Bernard, C., Lalonde, R., & Rebai, M. (2008). A prefrontal ERP involved in decision making during visual duration and size discrimination tasks. International Journal of Neuroscience, 118(1), 149–162.

4. Coull, J. T., Charras, P., Donadieu, M., Droit-Volet, S., & Vidal, F. (2015). SMA selectively codes the active accumulation of temporal, not spatial, magnitude. Journal of Cognitive Neuroscience, 27(11), 2281–2298.

5. Schlichting, N., de Jong, R., & van Rijn, H. (2020). Performance-informed EEG analysis reveals mixed evidence for EEG signatures unique to the processing of time. Psychological research, 84(2), 352–369.

6. Kononowicz, T. W., & van Rijn, H. (2014). Decoupling interval timing and climbing neural activity: a dissociation between CNV and N1P2 amplitudes. Journal of Neuroscience, 34(8), 2931–2939.

7. Bannier, D., Wearden, J., Le Dantec, C. C., & Rebaï, M. (2019). Differences in the temporal processing between identification and categorization of durations: a behavioral and ERP study. Behavioural brain research, 356, 197–203.

8. Wiener, M., & Thompson, J. C. (2015). Repetition enhancement and memory effects for duration. Neuroimage, 113, 268–278.

9. Paul, I., Le Dantec, C., Bernard, C., Lalonde, R., & Rebaï, M. (2003). Event-related potentials in the frontal lobe during performance of a visual duration discrimination task. Journal of clinical neurophysiology, 20(5), 351–360.

10. Lindbergh, C. A., & Kieffaber, P. D. (2013). The neural correlates of temporal judgments in the duration bisection task. Neuropsychologia, 51(2), 191–196. doi:10.1016/j.neuropsychologia.2012.09.0.

11. Paul, I., Wearden, J., Bannier, D., Gontier, E., Le Dantec, C., & Rebaï, M. (2011). Making decisions about time: Event-related potentials and judgements about the equality of durations. Biological Psychology, 88(1), 94–103.

12. Gontier, E., Le Dantec, C., Leleu, A., Paul, I., Charvin, H., Bernard, C., Lalonde, R., & Rebaï, M. (2007). Frontal and parietal ERPs associated with duration discriminations with or without task interference. Brain research, 1170, 79–89.

13. Gontier, E., Paul, I., Le Dantec, C., Pouthas, V., Jean-Marie, G., Bernard, C., Lalonde, R., & Rebaï, M. (2009). ERPs in anterior and posterior regions associated with duration and size discriminations. Neuropsychology, 23(5), 668.Gontier et al. 2009;

14. Tarantino, V., Ehlis, A. C., Baehne, C., Boreatti-Huemmer, A., Jacob, C., Bisiacchi, P., & Fallgatter, A. J. (2010). The time course of temporal discrimination: an ERP study. Clinical Neurophysiology, 121(1), 43–52.

15. Walther, A., Nili, H., Ejaz, N., Alink, A., Kriegeskorte, N., & Diedrichsen, J. (2016). Reliability of dissimilarity measures for multi-voxel pattern analysis. Neuroimage, 137, 188–200.

16. Kriegeskorte, N., Mur, M., & Bandettini, P. A. (2008). Representational similarity analysis-connecting the branches of systems neuroscience. Frontiers in systems neuroscience, 2, 4.

17. Brainard, D. H. (1997). The psychophysics toolbox. Spatial vision, 10(4), 433–436.

18. de Gardelle and Summerfield (2011) de Gardelle, V., & Summerfield, C. (2011). Robust averaging during perceptual judgment. Proceedings of the National Academy of Sciences, 108(32), 13341–13346.

19. Oostenveld, R., Fries, P., Maris, E., & Schoffelen, J. M. (2011). FieldTrip: open source software for advanced analysis of MEG, EEG, and invasive electrophysiological data. Computational intelligence and neuroscience, 2011.

20. Chaumon, M., Bishop, D. V., & Busch, N. A. (2015). A practical guide to the selection of independent components of the electroencephalogram for artifact correction. Journal of neuroscience methods, 250, 47–63.

21. Wichmann, F. a. & Hill, N. J. The psychometric function: I. Fitting, sampling, and goodness of fit. Perception & psychophysics 63, 1293–313 (2001).

22. Prins, N & Kingdom, F. A. A. (2018). Applying the Model-Comparison Approach to Test Specific Research Hypotheses in Psychophysical Research Using the Palamedes Toolbox. Frontiers in Psychology, 9:1250. doi: 10.3389/fpsyg.2018.01250.

23. JASP Team (2020). JASP (Version 0.12.2) [Computer software].

24. Wolff, M. J., Ding, J., Myers, N. E., & Stokes, M. G. (2015). Revealing hidden states in visual working memory using electroencephalography. Frontiers in Systems Neuroscience, 9, 123.

25. Ledoit, O., & Wolf, M. (2004). Honey, I shrunk the sample covariance matrix. The Journal of Portfolio Management, 30(4), 110–119.

26. Groppe, D. M., Urbach, T. P., & Kutas, M. (2011). Mass univariate analysis of event‐related brain potentials/fields I: A critical tutorial review. Psychophysiology, 48(12), 1711–1725.

27. Tjur, T. (2009). Coefficients of determination in logistic regression models—a new proposal: The coefficient of discrimination. The American Statistician, 63(4):366–372.

28. O’connell, R. G., Dockree, P. M., & Kelly, S. P. (2012). A supramodal accumulation-to-bound signal that determines perceptual decisions in humans. Nature neuroscience, 15(12), 1729.

29. Twomey, D. M., Murphy, P. R., Kelly, S. P., & O’Connell, R. G. (2015). The classic P300 encodes a build‐to‐threshold decision variable. European journal of neuroscience, 42(1), 1636–1643.

30. Wiener, M., Parikh, A., Krakow, A., & Coslett, H. B. (2018). An intrinsic role of beta oscillations in memory for time estimation. Scientific reports, 8(1), 1–17.

31. Balcı, F., & Simen, P. (2014). Decision processes in temporal discrimination. Acta psychologica, 149, 157–168.

32. Balcı, F. and Simen, P. (2016). A decision model of timing. Current Opinion in Behavioral. Sciences, 8:94–101.

33. Simen, P., Balci, F., deSouza, L., Cohen, J. D., & Holmes, P. (2011). A model of interval timing by neural integration. Journal of Neuroscience, 31(25), 9238–9253.

